# A multi-partner symbiotic community inhabits the emerging pest *Pentastiridius leporinus*

**DOI:** 10.1101/2025.06.03.657594

**Authors:** Heiko Vogel, Benjamin Weiss, Fortesa Rama, Andre Rinklef, Tobias Engl, Martin Kaltenpoth, Andreas Vilcinskas

## Abstract

The planthopper *Pentastiridius leporinus* has emerged as a severe crop pest, rapidly expanding both its host plant range and the affected areas in central Europe. Originating as a monophagous herbivore of reed grass, *P. leporinus* recently adopted polyphagous feeding and is now a pest of sugar beet, potato, carrot, and onion, suggesting rapid ecological niche expansion. *P. leporinus* vectors two bacterial pathogens, the γ-proteobacterium *Candidatus* Arsenophonus phytopathogenicus (CAP) and the stolbur phytoplasma *Candidatus* Phytoplasma solani (CPS), which are responsible for a range of disease syndromes, including syndrome basses richesses (SBR) in sugar beet. We used long-read metagenomic sequencing to characterize the genomes of microbes associated with *P. leporinus*, resulting in the complete sequences of CAP and CPS, as well as primary symbionts of the genera *Purcelliella, Sulcia* and *Vidania*, and facultative symbionts *Rickettsia* and *Wolbachia*. The primary symbionts are inferred to provide all ten essential amino acids and contribute to B vitamin biosynthesis. The genomes of CPS and CAP encode numerous pathogenicity factors, enabling the colonization of different hosts. Bacterial fluorescence *in situ* hybridization revealed the tissue distribution, cellular localization, relative abundance and transmission patterns of these bacteria. The intracellular presence of all primary symbionts in bacteriomes, the intracellular presence of *Wolbachia*, and the intranuclear localization of *Rickettsia*, suggest vertical transmission. CPS was restricted to salivary glands, suggesting strict horizontal, plant-mediated transmission, whereas CAP colonized all tissue types, allowing for horizontal and vertical transmission. Our data suggest that *P. leporinus* hosts an exceptionally broad range of symbionts, encompassing mutualistic, commensal and pathogenic interactions.

**Importance:** The planthopper *Pentastiridius leporinus* has recently expanded its host plant range and emerged as severe pest of sugar beet and potato crops in central Europe, which is exacerbated by its capacity to vector bacterial pathogens to its host plants. Because microbial symbionts may play an important role for both the host plant shifts and the transmission of pathogens, we used metagenomic sequencing and fluorescence *in situ* hybridization to characterize the microbial community associated with *P. leporinus*. We detected three bacteriome-localized primary symbionts that together provision all essential amino acids and several B-vitamins to the host, as well as two intracellular bacteria with a broader tissue distribution. In addition, we infer localization, transmission, and putative pathogenicity factors for the two major phytopathogens that are vectored by *P. leporinus*. Our results reveal a complex community of symbiotic bacteria that likely shapes the interaction of this emerging pest with its host plants.

## Introduction

Sap-feeding insects of the order Hemiptera are usually associated with symbionts that provide essential amino acids and vitamins (1-5). Some of these primary bacterial symbionts have been maintained by their hosts for hundreds of millions of years, whereas others have been replaced or complemented more recently by co-obligate symbionts (3,5). These more recently acquired symbionts are recruited from different bacterial groups, such as *Pantoea-Erwinia*, and can include not only insect-associated species but also plant pathogens and environmental bacteria that provide amino acids and vitamins, as well as other metabolic supplements (4). Hemiptera also frequently harbor and vector plant pathogens, especially viruses and bacteria, contributing to significant crop losses and economic impact (6).

*Pentastiridius leporinus* (Linnaeus, 1761, Hemiptera: Cixiidae) (Fig. 1A) belongs to the suborder Auchenorrhyncha (planthoppers, leafhoppers, spittlebugs, and cicadas), which often harbor diverse communities of bacterial endosymbionts (7,8). Planthoppers may have coevolved in an ancient relationship with bacterial symbionts of the genera *Sulcia* and *Vidania*, suggesting a multi-partner heritable community with different strategies to ensure vertical transmission (9). *P. leporinus* was originally reported to feed on reed grass (*Phragmites australis*; Poaceae) and is found on the endangered species list in Germany. Recently, however, it has rapidly expanded both its geographical distribution and host range, including crops such as sugar beet (*Beta vulgaris* subsp. *vulgaris*; Amaranthaceae) (10), potato (*Solanum tuberosum*; Solanaceae) (11-13), onion (*Allium cepa*; Amaryllidaceae) (14), winter wheat (*Triticum aestivum*; Poaceae), spring barley (*Hordeum vulgare*; Poaceae), and carrot (*Daucus carota* subsp. *sativus*; Apiaceae). *P. leporinus* and other planthoppers feed primarily on phloem sap, not only causing direct host plant damage but also enabling the vectoring of plant pathogens. Accordingly, *P. leporinus* is a major vector of ‘syndrome basses richesses’ (SBR), which causes severe economic losses in sugar beet and potato crops (15,16), but classical pest management has failed (17). SBR is caused by two phloem-restricted bacterial pathogens, the γ-proteobacterium *Candidatus* Arsenophonus phytopathogenicus (CAP) and the stolbur phytoplasma *Candidatus* Phytoplasma solani (CPS) of the stolbur group (16SrXII). Phytoplasmas are small bacteria lacking a cell wall, can infect many plant species, multiply within phloem-feeding insects of the Hemiptera order operating as transmission vectors, and can manipulate its host plants by modulating host defense and morphogenesis (18-20). Both phytopathogens, CAP and CPS are prevalent in sugar beets and potatoes displaying SBR-symptoms and are vectored by *P. leporinus* (12,13). SBR symptoms include the yellowing of older leaves, lancet-shaped leaf deformations, and necrosis of the vascular tissue, reducing beet sugar yields by up to 25% (21). The spreading of SBR in Germany is thought to be driven by climate change (16).

**Fig. 1.**
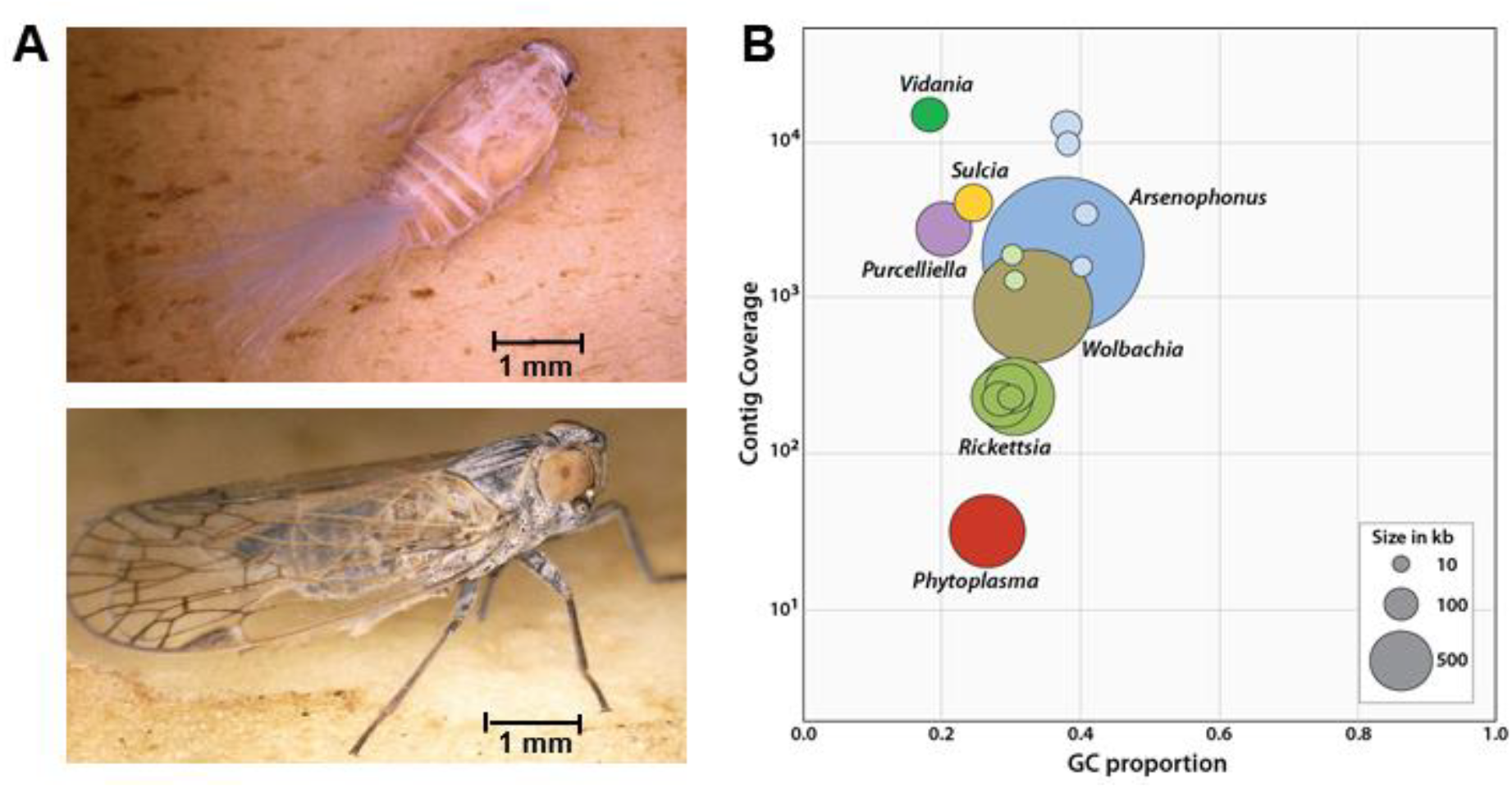
The planthopper *Pentastiridius leporinus* and its associated bacterial symbionts and pathogens. (A) *P. leporinus* nymph (top) and adult (bottom). (B) Visual representation of global genome features of the endosymbionts and pathogens associated with *P. leporinus* based on metagenomic assemblies. Taxon-annotated GC proportion-coverage plots are based on the final, short-read-polished long-read assemblies without the *P. leporinus* genomic contigs. Colored circles represent contigs or complete circular chromosomes and the lighter variants of the same colors represent plasmids. The *x*-axis shows the GC proportion and the *y*-axis shows sequence coverage for all contigs.

The rapid ecological niche expansion of *P. leporinus* requires adaptations to the diverse nutritional and phytochemical compositions of distantly related crops. Vertically transferred bacterial symbionts play a key role in the evolution of sap-feeding insects not only by providing nutrients but also by conferring adaptations to biotic and abiotic stress (22-24). However, primary symbionts with highly eroded genomes are unlikely to mediate adaptations to new host plants or expand the geographic distribution of their insect hosts, so new symbionts may be responsible for this phenomenon (5, 25, 26). CAP and CPS are vector-borne cross-kingdom pathogens spanning the parasitism–mutualism continuum (28, 29) and are able to replicate in plant sieve elements and in hemipteran vectors. It remains unclear whether *Arsenophonus* spp. were originally plant pathogens that adapted to sap-sucking insects to facilitate transmission, or insect symbionts that became pathogenic in plants. Comparative genome analysis supports the “insect first” scenario (29), but *Arsenophonus* also shifts along the parasitism– mutualism continuum and may supplement the nutritionally poor diet of its sap-feeding hosts with B vitamins and essential amino acids in order to increase host fitness and promote transmission from infected to uninfected host plants (30). Accordingly, CAP and CPS may be both plant pathogens and insect symbionts, but their effect on host fitness is insufficiently understood. Here, we characterized bacterial symbionts associated with *P. leporinus* in the context of its rapid host range expansion, seeking to understand their ability to benefit the insect host and encode virulence factors that facilitate the colonization of insects and/or plants. We also used fluorescence *in situ* hybridization (FISH) to determine the tissue distribution of symbionts, providing insight into their transmission routes.

## Results and discussion

### Genomes of *P. leporinus* obligate nutritional endosymbionts

We cataloged the bacteria in two laboratory specimens (one female and one male) and one field specimen of *P. leporinus* by combining single-molecule long-read and Illumina short-read sequencing. Each metagenomics dataset resulted from several assemblies, including read subsampling, sequences with >10-kb read length cut-offs and bacteria-only datasets. Our untargeted screening revealed a complex symbiont community, including CAP, CPS and bacteria of the genera *Sulcia, Vidania, Purcelliella, Rickettsia* and *Wolbachia* (Table 1, Fig. 1B). The assemblies of the three *P. leporinus* individuals showed variability in the presence or absence of certain bacteria. CAP and CPS were absent from the female laboratory specimen whereas CPS was absent from the male but CAP was abundant. However, subsequent qPCR-based screening of several laboratory-reared individuals confirmed the presence of CAP and CPS in most adults (data not shown). Few reads mapped to the assembled CAP and CPS genomes because the bacterial titers were sometimes too low for successful *de novo* genome assemblies. In contrast, the field-collected specimen carried all seven bacterial taxa, including CAP and CPS.

**Table 1:**
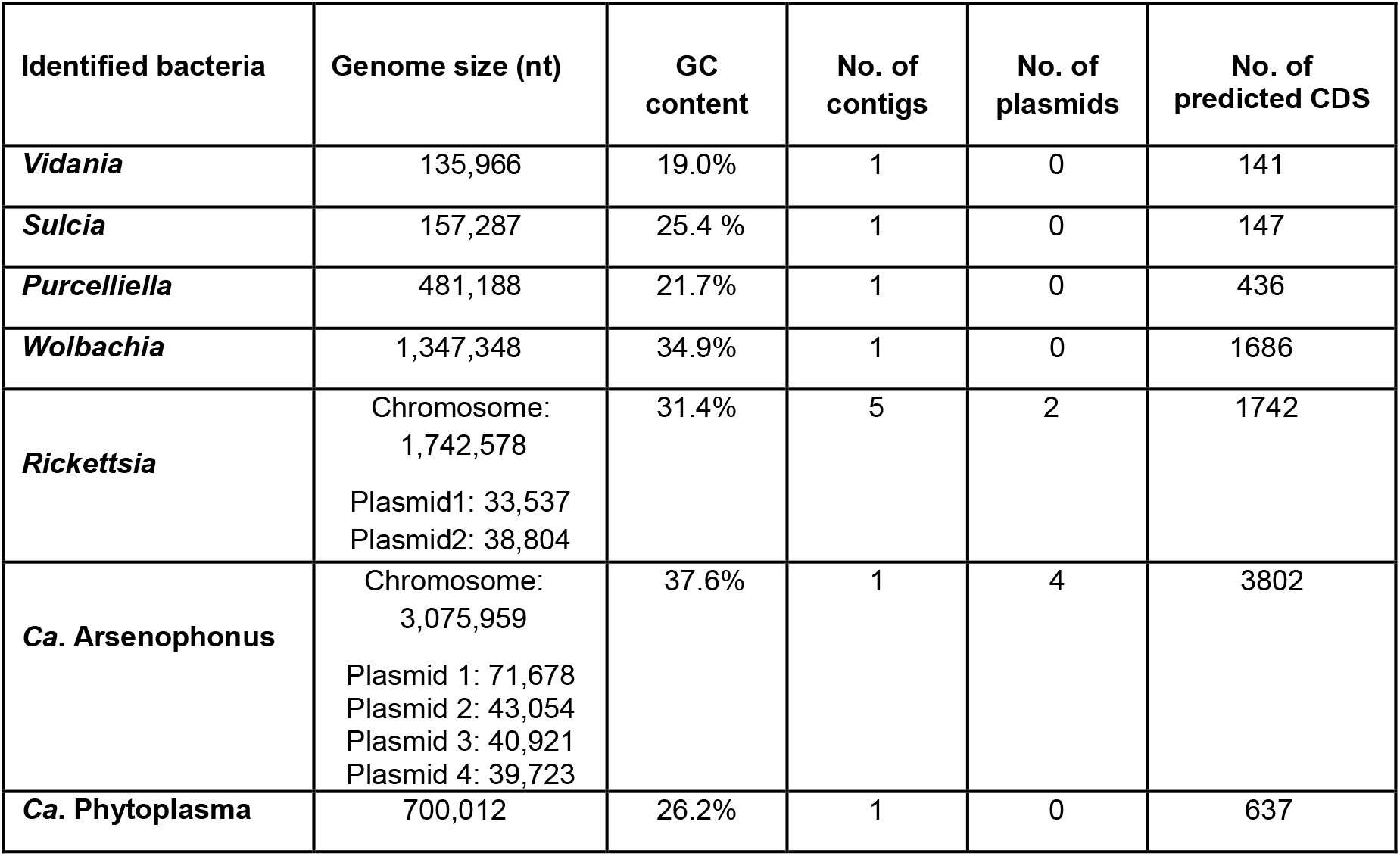
Results on endosymbiont genome assemblies and gene predictions.

The complexity of the symbiont community in *P. leporinus* has been reported in other members of the hemipteran suborder Auchenorrhyncha, in which nutritional symbiosis was established ∼300 Mya (4,7,8). The coevolution of ancient symbionts with their phloem-feeding insect hosts allowed genome reduction, but not all bacterial associations in Auchenorrhyncha are static. Some ancient symbionts have been replaced or complemented by recently acquired bacteria whose genomes have also undergone subsequent erosion. The gene-dense genomes of ancient planthopper symbionts *Sulcia* (∼157 kb) and *Vidana* (∼135 kb) show much more extensive erosion than the more recently acquired *Purcelliella* (∼481 kb) (Fig. 2A). All three primary symbionts were found to be very AT-rich and abundant in all three *P. leporinus* specimens (Fig. 1B).

**Fig. 2.**
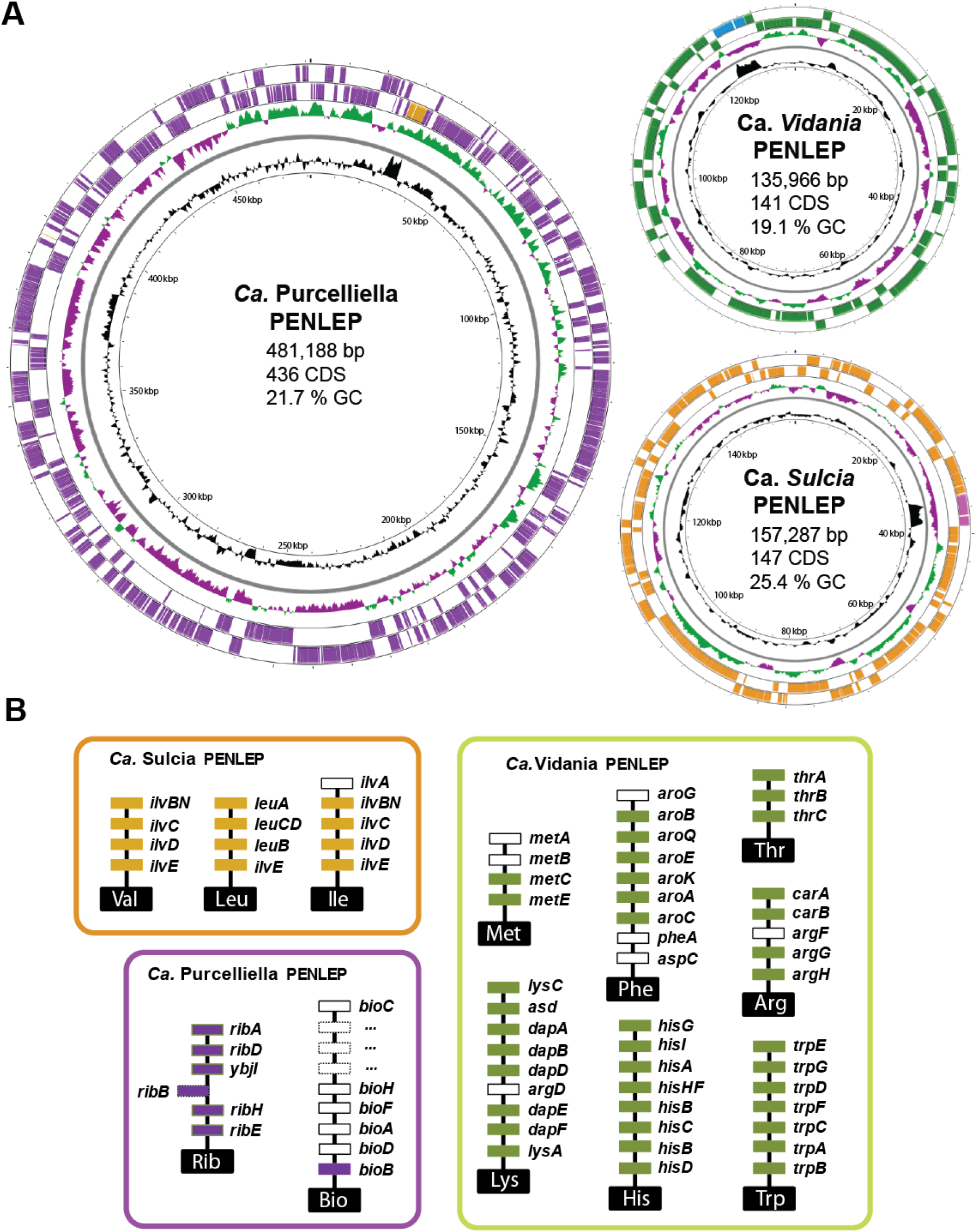
Genome maps and major biosynthetic pathways of *P. leporinus* primary symbionts. (A) Visualization of circular genomes and basic genome characteristics of the three primary symbionts *Ca*. Vidania, *Ca*. Sulcia and *Ca*. Purcelliella identified in the metagenomic assemblies. Genes on the forward and reverse strands are shown in color and the rRNA operon is shown in a contrasting color. The innermost circle shows the genome size, the second shows the GC content distribution, and the third shows the relative GC skew. Additional genomic information, including the number of coding sequences (CDS), is shown inside each genome circle. (B) Symbiont-encoded amino acid and B vitamin synthesis pathways contributing to *P. leporinus* metabolism. Genes encoding enzymes in the pathways predicted from bacterial genome sequences are shown in solid blocks and genes that could not be identified are shown in white rectangles. Abbreviations: Bio = biotin, His = histidine, Ile = isoleucine, Leu = leucine, Lys = lysine, Met = methionine, Phe = phenylalanine, Rib = riboflavin, Thr = threonine, Trp = tryptophan, Val = valine.

In several cixiid planthoppers, *Vidania* produces seven essential amino acids, whereas *Sulcia* produces only the branched-chain amino acids valine (Val), leucine (Leu) and isoleucine (Ile) (4,9). We found that the *Sulcia* genome isolated from *P. leporinus* encoded all enzymes needed for the synthesis of Val and Leu, and all but one needed for the synthesis of Ile (Table S1). The recovered *Vidania* genome also has complete sets of genes for histidine (His), tryptophan (Trp) and threonine (Thr) synthesis, but one or more was missing from the methionine (Met), phenylalanine (Phe), lysine (Lys) and arginine (Arg) pathways (Fig. 2B, Table S2). In the case of Arg, our recovered *Vidania* sequence lacked only *argF*, although this gene was present in *Vidania* isolated from three fulgorid planthoppers (31). Most of the gene losses we identified have been reported previously in *Sulcia* and *Vidania* (4). The last step (*pheA* and *aspC*) in the phenylalanine biosynthesis and the cysteine pathway (*metAB*) in methionine biosynthesis of *Vidania*, and in *Sulcia* the initial step (*ilvA*) in isoleucine biosynthesis could not be identified.

In the cixiid *Oliarus filicicola*, the more recently acquired γ-proteobacterial symbiont *Purcelliella* contributes the synthesis of B vitamins and probably cysteine (Cys) (4, 32). In *P. leporinus*, the *Purcelliella* symbiont genome encoded all enzymes in the riboflavin (Rib) pathway, but only a single gene in the biotin (Bio) pathway (Fig. 2B, Table S3). Missing annotations due to A/T-rich genomes are unlikely to explain the absence of genes because the coding density in small bacterial genomes is high, and at least the amino acid biosynthesis genes are generally well annotated. It is unclear how these pathways are completed to provide nutrition, but it is likely that bifunctional or substrate-promiscuous enzymes are involved in the missing steps or, for terminal aminotransferases, they are complemented by the host (32-34). An alternative explanation that needs to be further investigated is the occurrence of horizontal gene transfer events from bacterial donors to the insect host genome. The acquisition of certain vitamins from plant sap has also been proposed (35,36). The genome erosion of *Sulcia, Vidania* and *Purcelliella* should also result in a demand for metabolic resources from the host in order to maintain these beneficial relationships. In the leafhopper *Macrosteles quadrilineatus*, and several psyllid species, the host expresses genes in the bacteriocytes that may complement the loss of genes required for cell wall modifications, nucleic acid synthesis, DNA repair, transcription and translation, allowing the bacteria to provide essential nutrients (37,38).

### Genomes of *Rickettsia* and *Wolbachia*

*Rickettsia* is a genus of Gram-negative bacteria that can be insect pathogens or symbionts. Due to their obligate intracellular lifecycle, they have tools for the invasion of host cells and subsequent evasion of immune responses, including effectors and membrane proteins involved in cytoskeletal rearrangement and adhesion (40-42). Like other *Rickettsia* spp., the genome of *Rickettsia* associated with *P. leporinus* encodes a full Sec-dependent pathway as well as a type IV secretion system (T4SS). We identified 100 genes with Sec signal peptide (SP) sequences, including homologs of *sca2, ompB*, several putative adhesin and porin family genes, as well as genes encoding virulence factors such as ecotin (Table S4). We also found genes encoding a number of candidate T4SS effectors such as RalF, which is reportedly involved in the phagocytosis of *R. typhi* in mammalian cell lines, as well as several ankyrin-repeat proteins previously shown to prevent phagosome fusion with host endosomes, thus aiding in evading endocytic maturation (40,43). We also identified *sca4*, encoding a multifunctional effector involved in the adhesion and invasion of host cells (41).

Similar to *Rickettsia*, the genus *Wolbachia* engages in a large variety of symbiotic interactions with arthropods, ranging from entomopathogenic to obligately mutualistic. Our assembled *Wolbachia* genome encoded Sec and twin-arginine translocation (TAT)-dependent protein translocation pathways, as well as the T4SS typically found in insect-associated strains (39). 52 genes are predicted to include a Sec-SP sequence, most of which encode hypothetical proteins. We did not identify any potential effectors in the predicted list of genes containing a Sec-SP sequence (Table S5). Endosymbiotic *Wolbachia* species are also known to induce several reproductive phenotypes in their insect hosts, including cytoplasmic incompatibility (CI), parthenogenesis and male killing. We identified homologues of the CI factors *cifA* and *cifB*, which regulate cytoplasmic incompatibility to manipulate host reproduction, resulting in embryonic lethality, but also a homolog of the *wmk* gene involved in *Wolbachia*-mediated male killing in *Drosophila*.

However, *Rickettsia* and *Wolbachia* can also be mutualistic endosymbionts, contributing vitamins and other nutrients, boosting the fitness of its insect hosts (44, 45). Indeed, *Wolbachia* encodes the full biotin and riboflavin biosynthetic pathways, potentially supplementing these B vitamins to *P. leporinus*. Since *Purcelliella* only encodes the last step of biotin production, *Wolbachia* could potentially surrogate the role of a biotin producing endosymbiont. It is so far unclear if *Rickettsia* provides *P. leporinus* with any nutrients.

### Genomes and candidate pathogenicity factors of vectored plant pathogens

#### *Ca*. Phytoplasma

The genome of CPS from a different population of *P. leporinus* has recently been published (46) and our independent analysis confirmed a similar ∼700 kb circular chromosome with 637 predicted coding sequences (Table 1, Fig. 1, Fig. 3A, Table S7). Our CPS genome was predicted to encode a complete Sec pathway, which can secrete bacterial effectors. We also found 23 genes containing a Sec-SP, 11 representing the sequence variable mosaic (SVM) family. Several validated phytoplasma effectors and many candidates contain an SVM domain, including SAP54/PHYL1, which is released into the phloem and regulates plant development at the level of transcription (47) (Fig. 3B). However, only partial hits for SAP54 (GenBank: MK858224.1) were found, as was the case for the other well-characterized *Phytoplasma* effectors SAP11 (GenBank: CP000061.1 “AYWB_370”) and TENGU (GenBank: AB750357). The predicted CPS proteins showed low levels of sequence identity to SAP54, SAP11 and SAP05, suggesting they are homologs but not orthologs (although see 46). Even so, other SAP orthologs have been identified in CPS, such as SAP09-like, SAP45-like, SAP50-like, and SAP61-like. These show a higher level of sequence identity across different *Phytoplasma* strains and species and are largely uncharacterized, although some predicted proteins seem to include docking sites for plant cell cycle regulators (48). We also identified hemolysin-related proteins (HRPs) with Sec-SP sequences. These have been linked to pore-forming activity and virulence in other Mollicutes, but their precise function is unknown (49). Our CPS genome also encoded co-chaperonins GroES and GroEL (ORF948/950), forming a protein folding system that can trigger plant defenses during insect herbivory and play a role in symbiotic microbe–insect interactions (51).

**Fig. 3.**
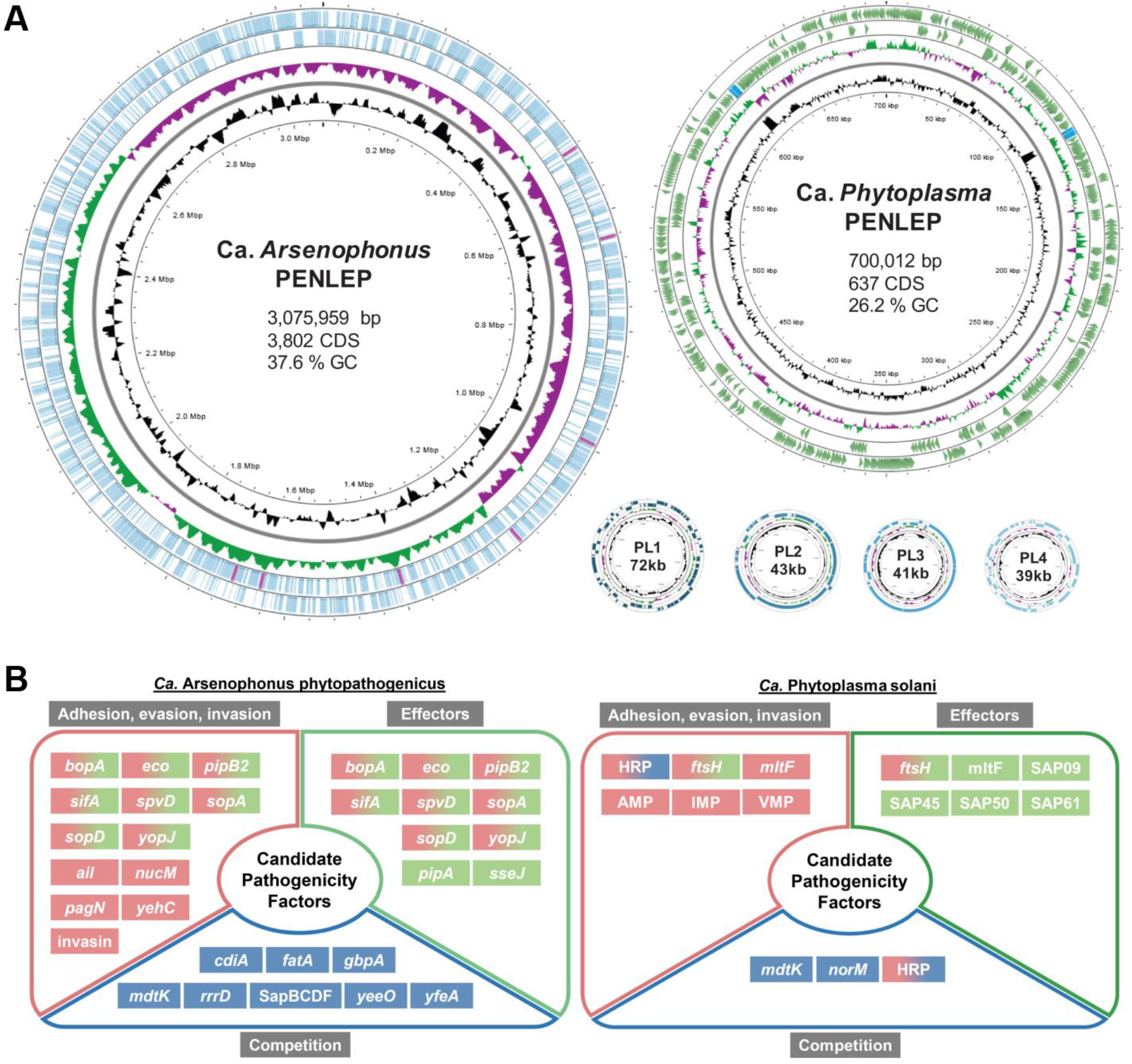
Genome maps and predicted candidate effectors of *P. leporinus*-transmitted plant pathogens. (A) Visualization of the circular genomes and basic genome characteristics of the vectored plant pathogens Ca. *Arsenophonus phytopathogenicus* and Ca. *Phytoplasma solani* in the metagenomic assemblies. PL1-4 depict Ca. *Arsenophonus* plasmids. Genes on the forward and reverse strands are shown in color and the rRNA operon is shown in a contrasting color. The innermost circle shows the genome size, the second shows the GC content distribution, and the third shows the relative GC skew. Additional genomic information, including the number of coding sequences (CDS), is shown inside each genome circle. (B) Candidate pathogenicity-related factors of *Ca*. Arsenophonus and *Ca*. Phytoplasma based on genome and gene prediction analysis. Abbreviations: AMP = antigenic membrane protein, HRP = hemolysin-related protein, IMP = immunodominant membrane protein, SAP = secreted Aster Yellows Witches’ Broom protein, VMP = variable membrane protein.

Phytoplasmas lack a cell wall, so all membrane proteins are in direct contact with the environment. Antigenic membrane proteins (AMPs), immunodominant membrane proteins (IMPs) and variable membrane proteins (VMPs) are promising candidate pathogenicity and symbiosis-related factors that have been confirmed as such in Flavescence dorée phytoplasma (FDP) and *Ca*. P. mali (52,53). In our *P. leporinus*-associated *Phytoplasma* genome, we found six genes encoding these membrane proteins, comprising one AMP, one IMP, and four VMPs.

Membrane-bound proteases can also act as pathogenicity factors. FtsH (HflB) is one of five AAA+ proteases in *Escherichia coli*, but is the only one essential for cell viability (54). These ATP-dependent zinc proteases are involved in membrane protein quality control, including the removal of inactive membrane-bound Sec complexes (55). In *E. coli* and most other prokaryotes, including Mollicutes such as mycoplasmas and spiroplasmas, *ftsH* is a single-copy gene (56), whereas several copies are often found in phytoplasmas. In *Ca*. P. mali and *Ca*. P. vitis, FtsH is also a putative virulence effector (57,58). The CPS genome contains 15 copies of *ftsH* (ftsH1–15), with *ftsH3, ftsH12* and *ftsH15* predicted to include a Sec-SP. InterPro domain prediction showed a variety of membrane-spanning topologies, typically one or two transmembrane domains and a large non-cytoplasmic domain that includes a metalloprotease and AAA+ ATPase. The *ftsH* homologs are structurally diverse, suggesting equally diverse functions. In *Ca*. P. vitis, the expression profiles of *ftsH* paralogs differed between insect and plant hosts, suggesting a dynamic role during infection and colonization (57).

#### *Ca*. Arsenophonus

The sequencing and assembly of the complete CAP genome resulted in a single contig with a chromosome size of 3.08 Mb (containing 3,802 predicted coding sequences; Table 1, Fig. 3A, Table S7) and four plasmids ranging in size from 39 to 72 kb. A comparative analysis of *Arsenophonus* genomes showed a large size range, from ∼663 kb to ∼4.9 Mb (30). The recently published CAP assemblies from *P. leporinus* populations in Switzerland and France show similar sizes to the genome presented in this study (29). Many *Arsenophonus* species, including CAP, encode a full complement type III secretion system (T3SS), a Sec-dependent pathway, and a TAT pathway (Fig. 3B). In many bacteria, T3SS is needed for motility and virulence, and proteins secreted by T3SS are well known effectors involved in a wide range of diseases (60). We also found several T3SS effector genes, some reportedly involved in the invasion of host tissues (*bopA, pipB2, sopD* and *sifA*) and cells (*sapB*). Furthermore, we identified homologs of the *Salmonella* genes encoding virulence factors PipA, SpvD and SseJ, which inhibit host inflammatory responses (59), and putative toxins of the repeats-in-toxin (RTX) exoprotein, colicin V, heat-labile enterotoxin, insecticidal toxin complex (SidC), TolA, and OmpA families (61).

We found 223 candidate genes encoding effector proteins with predicted SPs (218 with a Sec-SP and five with a TAT-SP), some of which can be classified as virulence factors or putative effectors. They included adhesion proteins such as PagN, Ail, and YehC, as well as other virulence factors such as YopJ, ecotin and NucM. We identified 12 *nucM* homologs, encoding non-cytoplasmic endonucleases that evade host immune responses by degrading nucleic-acid extracellular traps (62). The remaining SP-containing genes of interest encoded siderophores (*fatA, yfeA*), lysozymes (*rrrD)* and lytic monooxygenases (*gbpA*). We found other adhesion/invasion-related genes lacking typical SP sequences, one encoding an invasin domain-3 protein that binds eukaryotic integrin receptors to trigger bacterial uptake by host cells (63).

Toxin-antitoxin (TA) systems encoded by the CAP genome included the contact-dependent growth inhibition system involving CdiA, which contains a SP and can confer an advantage in inter-bacterial competition (64). Several multidrug and toxic compound extrusion (MATE) family transporter genes were identified, such as *mdtK* and *yeeO*, which are involved in antimicrobial resistance (65). Genes were present for all parts of the multicomponent polyamine exporter SapBCDF, which contributes to resistance against antimicrobial peptides in some enterobacteria (66), suggesting that CAP can maintain cell viability in competitive microbial environments.

The recently published analysis of CAP genomes from *P. leporinus* populations in France and Switzerland showed a relatively high percentage of phage-derived regions (29). We identified the chromosome-encoded phage resistance genes *abiGI*, which is part of the lactococcal abortive infection system, as well as a plasmid-encoded cyclic oligonucleotide-based antiphage signaling system III (CBASS III), both of which are versatile antiphage defense systems in a number of bacterial species (67,69,70). We also identified a type I bacterial restriction-modification (RM) system encoded on the chromosome. Additionally, a plasmid-encoded RM Type IIG system can be found, as well as homologs of another RM Type II system. We saw no evidence of a chromosome-encoded CRISPR/Cas system, but several pseudogenes of a CRISPR-associated endo-deoxyribonuclease were present on plasmids 1 and 4, with no CRISPR sequences identified on either plasmid or on the chromosome. The CRISPR/Cas system is often highly pseudogenized or lost in obligate *Arsenophonus* endosymbionts, as are other phage defense systems (71). This loss has been linked to genome expansion through mobile genetic elements, as well as the transmission mode, with horizontally transferred strains having more diverse anti-phage systems than strains that are vertically transmitted or experience a mixture of vertical and horizontal transmission (71). However, CAP maintained a number of phage defense systems, despite likely being vertically and horizontally transmitted, thus not fitting the adaptation model proposed in (71). Other plasmid-borne genes encoded an F-like type IV secretion system, lysozymes, and toxin-antitoxin systems associated with reproductive parasitism (72) and could thus be relevant for the wide distribution of CAP in *P. leporinus* tissues.

### Tissue distribution of *Pentastiridius*-associated microorganisms

We used our *P. leporinus* metagenomics data to design species-specific FISH probes and map the distribution of microbes in males, females and nymphs randomly selected from our laboratory population or based on a qPCR prescreening to identify individuals with and without CAP and/or CPS. The analysis of multiple tissue sections and samples revealed a complex array of bacteriomes but also unknown structures (Fig. 4A, Fig. S1). Based on the morphological data, we differentiated two types of salivary glands (Fig. 4A, I & II) and four types of bacteriomes (Fig. 4A, structures III-VI, Fig. 4C-K). Type I-II structures (Fig. 4A-C) are probably salivary glands adjacent to the intestine, characterized by low bacterial density. The type III bacteriome of females displayed a high density of bacteria, and the typical irregular structure of bacterial cells with eroded genomes (Fig. 4D, E; Fig. S1), but these fluorescent signals could not be clearly assigned to one of the bacterial species identified in our metagenomics data. It is highly unlikely that we have missed to identify an additional bacterial species in our metagenome data. Since the labeled eubacterial probe EUB-338-Cy7 provided a clear bacterial signal, peculiarities of type III bacteriome membrane structures could hinder the access of other fluorescent dyes and thus effective species-specific 16S labeling. Bean-shaped type IV bacteriomes displayed a high density of *Vidania* (Fig. 4F, G; Fig. S1), and the paired, elongated bacteriome V is tightly packed with *Purcelliella* primary symbiont cells (Fig. 4H, I; Fig. S1). The egg-shaped bacteriome VI shows a high density of *Sulcia* cells (Fig. 4J, K).

**Fig. 4.**
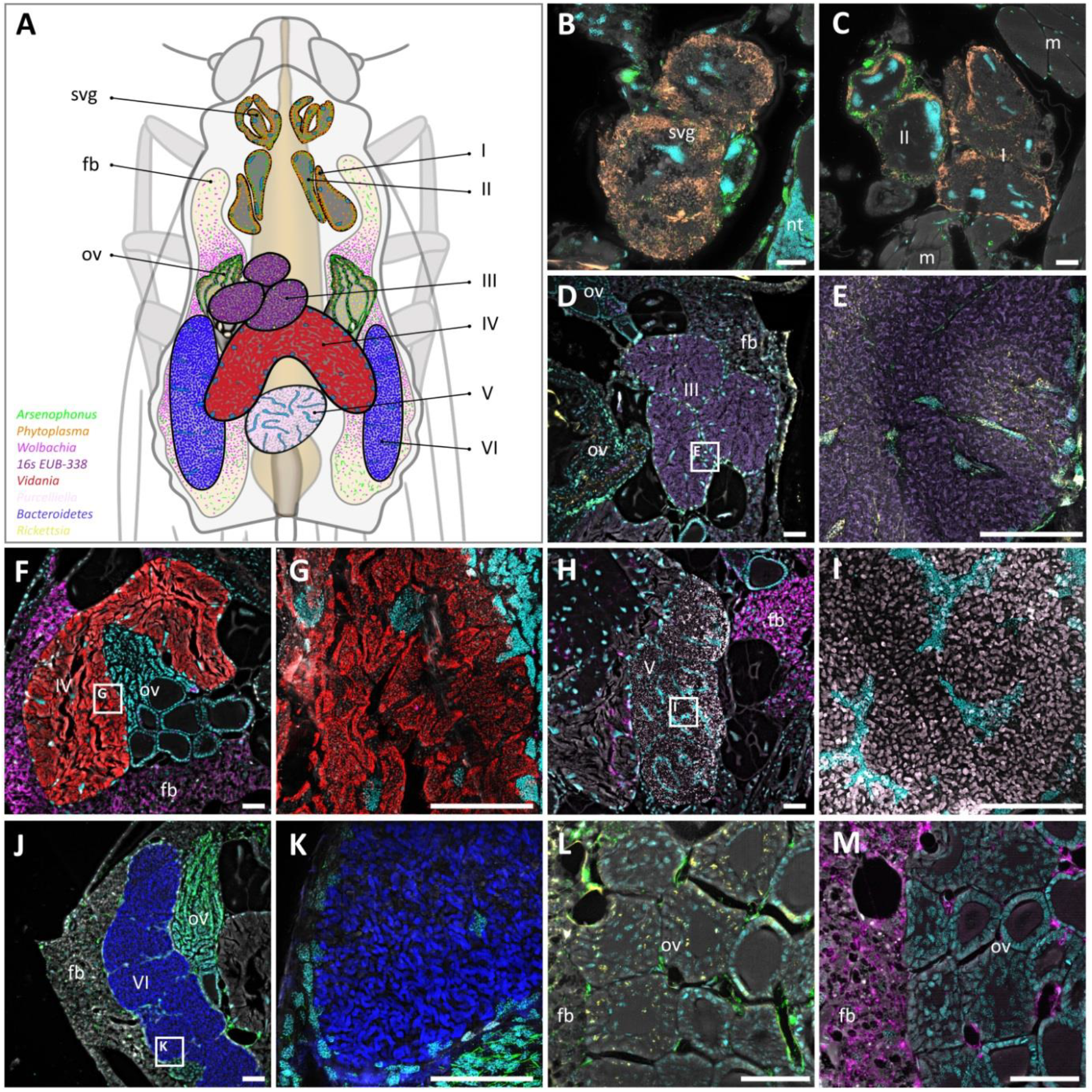
Localization of bacteria by fluorescence in situ hybridization (FISH) in semi-thin transverse sections of *Pentastiridius leporinus*. The labeled eubacterial probe EUB-338-Cy7 (magenta) was used with DAPI to counterstain the host nuclei (cyan). **A**. Schematic representation of the localization of organ structures identified in *P. leporinus*. Labels I-VI depict different tissue structures containing bacteria or bacteriomes. Svg = anterior salivary gland, fb = fat body, ov = ovary. **B**. *Phytoplasma* and *Arsenophonus* identified in salivary glands (svg). *Arsenophonus* can also be found in parts of the nervous system tissue (nt). **C**. *Phytoplasma* and *Arsenophonus* bacteria were localized in structures I and II (probably posterior salivary glands). *Arsenophonus* is also found in muscles (m). **D**. Structure III, filled with numerous unidentified bacteria (EUB-338), as well some *Rickettsia* and *Arsenophonus* in the ovary (ov). **E**. Detailed view of the structure III with unidentified bacteria (EUB-338), some *Arsenophonus*, and partly intranuclear *Rickettsia*. **F**. Structure IV, filled with a high density of *Vidania*. High densities of *Wolbachia* are also found in the fat body (fb) near the ovaries (ov). **G**. Detailed view of the structure IV with high density of *Vidania*. **H**. Structure V, filled with a high density of *Purcelliella*. A high density of *Wolbachia* is also found in the fat body (fb). **I**. Detailed view of structure V with high density of *Purcelliella*. **J**. Structure VI, filled with a high density of *Sulcia*. A high density of *Arsenophonus* is also found in the fat body (fb) and the ovary (ov). **K**. Detailed view of the structure VI with high density of *Sulcia*. **L**. Ovary (ov) with scattered cells of *Rickettsia* and *Arsenophonus* in the fat body (fb) as well between the ovaries. **M**. Ovary tissue with *Rickettsia* and *Arsenophonus* located in the nuclei (n). In panels B-M bacteria are colored according to the colors of the overview schematic shown in panel A. Scale bar = 50 µm.

We confirmed the presence of CAP in the salivary glands of *P. leporinus* supporting transmission from infected to uninfected host plants via phloem feeding as well as horizontal transmission among individuals feeding on the same host plant (15) (Fig. 4B, C). By contrast, *Arsenophonus* is likely also transmitted vertically because we found CAP in the fat body and endomysium, within bacteriomes, and in areas between the ovaries (Fig. 4L, M). CAP was also found in the epithelium surrounding the bacteriomes as well as between large bacteriome cells (Fig. 4D, E, J). Interestingly, *Arsenophonus* was also found inside cells. In contrast to the very broad tissue distribution of CAP, CPS was found exclusively in the salivary glands, co-occurring with CAP, whereas intranuclear *Rickettsia* were present in salivary glands and in the gut (Fig. 4B, C; Fig. S1).

The presence of CAP in many tissues (including ovaries; Fig. 4J, M), as well as its intracellular localization, indicates that CAP uses mixed-mode transmission in *P. leporinus*, as previously reported (15). The benefit of *Arsenophonus* for other insect hosts, such as the aphid *Aphis gossypii* and the corn planthopper *Peregrinus maidis* was explained by its ability to synthesize B vitamins and essential amino acids (18,24). *Arsenophonus* also interacts with other primary bacterial symbionts to synergistically compensate for unbalanced amino acid profiles in the phloem (24). However, *Arsenophonus* has negative effects in some insect species, such as the manipulation of host reproduction by male killing in *Nasonia* wasps (73). The numerous predicted effectors encoded in the CAP genome and plasmids may facilitate its establishment in host plants and widespread distribution in *P. leporinus* tissues. The acquisition of new symbionts with dual roles in the vector insect and its host plants may be an important driver of ecological niche expansion in sap-feeding insects (5) and was, for example, recently confirmed in mealybugs (74). Although the co-localization of CAP with other symbionts in *P. leporinus* could enable their interactions at various levels, this hypothesis needs experimental validation.

CPS can manipulate both its vector insect and host plant (20,75). The anticipated localization of CPS in *P. leporinus* salivary glands supports its transmission to sugar beet and potato plants via this vector. *Phytoplasma* infection renders host plants more susceptible to insect herbivores by making nutrients accessible and by depleting plant defensive compounds (76). For example, SAP11 triggers vector insect reproduction by influencing plant development and defense hormone biosynthesis (77). It is unclear whether CPS contributes to the broadening of the dietary niche of *P. leporinus*.

## Conclusions

In central Europe, the planthopper *P. leporinus* has recently shifted from a monophagous to a polyphagous lifestyle. *P. leporinus* is a sap-feeding insect that transmits at least two plant pathogens, so the shift has resulted in the emergence of a new insect pest causing severe economic damage in important crops. Although the expansion of the dietary niche may benefit *P. leporinus*, adaptations to the spectrum of nutritional compositions and chemical defenses in the expanding host plant range will likely incur costs. Symbiotic bacteria may drive ecological niche expansion and adaptive radiation in herbivorous insects (25,26) and could therefore play an important role in dietary shifts. Our analysis shows that *P. leporinus* uses its primary bacterial symbionts *Sulica, Vidania* and *Purcelliella* to compensate for nutritional deficits in its phloem-sap diet, such as all ten essential amino acids and several B vitamins. The roles of other symbiotic bacteria abundantly present in *P. leporinus*, i.e. *Wolbachia*, intranuclear *Rickettsia*, horizontally transmitted CPS localized in the salivary glands, and horizontally and vertically transmitted CAP localized in many different tissues currently remain speculative. However, the complex association of *P. leporinus* with seven bacterial symbionts provides an interesting system to investigate the influence of individual symbionts and their interactions on the expansion of the host’s ecological niche space and its capacity to vector pathogens to the host plant. Furthermore, insights into the role and interactions of microbial symbionts may provide novel leads for controlling this emerging and devastating agricultural pest insect.

## Materials and Methods

### Insect collection and rearing

Adult *P. leporinus* specimens were collected from sugar beet fields in South Hesse. Adults were kept in cages and were provided with ∼8-week-old sugar beets plants (Annarosa, KWS) in 3-L pots (Lamprecht-Verpackungen, Göttingen, Germany) covered with expanded clay. Eggs were transferred to boxes containing geohumus substrate (Geohumus, Frankfurt, Germany) and sugar beet plants. Nymphs and adults were kept at 22 °C and 60% relative humidity with a 16-h photoperiod. Detailed insect rearing conditions are described elsewhere (10).

### Metagenome sequencing and assembly

Genomic DNA was extracted from three *P. leporinus* specimens: one male and one female from our 2023 laboratory rearing population and one individual from a population collected in 2024 from a sugar beet field. Insect tissue was mixed with 500 µL CT buffer in a 2-mL microfuge tube and homogenized with ball bearings in a TissueLyser LT (Qiagen, Hilden, Germany). Genomic DNA was isolated using the Nanobind Big DNA kit (PacBio, Menlo Park, CA, USA) and high-molecular-weight (HMW) fragments were isolated using the Short Read Eliminator kit (PacBio). Isolated HMW DNA was cleaned using AMPure XP beads (Beckman Coulter, Krefeld, Germany). DNA purity and concentration were measured using a Qubit fluorimeter (Thermo Fisher Scientific, Darmstadt, Germany). DNA integrity was determined using a Tapestation 4150 (Agilent, Waldbronn, Germany).

End-DNA repair was carried out before Oxford Nanopore Technologies (ONT) library preparation using the NEBNext Ultra II DNA Library Prep Kit (New England Biolabs, Ipswich, MA USA). We added sequencing adapters using the V14 Ligation Sequencing Kit (ONT, Oxford, UK) and sequencing was carried out on a MinION Mk1B platform (ONT) using R10.4.1 flow cells (ONT). For each of the three samples, two flow cells were loaded with 70–100 ng of the libraries three times during a run of 96 h, with intervening washing steps using the Flow Cell Wash Kit (ONT). Super high-accuracy (SUP) base-calling of the raw reads with Guppy v6.0.1 (ONT) yielded 35–55 Gb of sequence data per specimen.

The three *P. leporinus* (meta)genome sequences were assembled separately using Flye v2.9.2 (78) in meta-mode, followed by four iterations of polishing using Racon, and one round of error correction using Medaka. For final error correction, we used ntEdit with paired-end (2 × 150-bp) Illumina data generated from the same HMW DNA as used for ONT sequencing. PurgeDups was used to remove duplications (heterozygous regions) and generate haploid genomes for downstream analysis. To optimize the assembly of insect-associated bacterial genomes, two additional assemblies per individual dataset were tested. First, all reads were classified against the RefSeq 2024-04 WGS database using Kraken2 (79) in OmicsBox v3.2, and reads assigned to bacteria were separated from all others. The bacteria-only reads were then assembled using Flye v2.9.2 with the “–meta” option. After polishing as above, taxon IDs were assigned to contigs using Blobtools. Second, reads were mapped to the bacterial assemblies using Geneious 2024.0.7 in high-sensitivity mode and three mapping iterations. The reads were assembled using Flye v2.9.2 and Unicycler v0.5.1 (80). All three assemblies per *P. leporinus* individual were compared and the optimal assembly selected based on criteria such as genome fragmentation and completeness (BUSCO) values.

### Finalizing and annotating bacterial genomes

Polypolish was used for final error correction primarily in repeat regions (79). Genome annotations including gene, phage-derived element and repeat predictions, as well as genome map visualizations were prepared using Prokka, Bakta (80), VirSorter, mobileOG-db (83), Phigaro (84) and Alien Hunter (https://www.sanger.ac.uk/tool/alien-hunter) on the Proksee server (85). Alternative gene predictions with Glimmer, BLASTX searches using the NCBI nr database (2024-07-11), conserved domain searches using InterProScan (86) and Gene Ontology (GO) classifications were applied using Omicsbox v3.2. KEGG pathway analysis was carried out using the server version of BlastKOALA (87). Potential virulence effector proteins encoded by the CAP and CPS genomes were predicted using the information obtained from SignalP 6.0 searches, InterPro and BLAST results obtained from the predicted gene annotations and BLAST searches against custom databases using known effectors. The web-based tool DefenseFinder was used to further identify anti-phage defense systems (88).

### Localization of bacterial symbionts and plant pathogens

The localization of symbionts and pathogens was assessed by FISH (89). Briefly, adults and nymphs (legs and, if present, wings removed) were fixed in 80% *tert*-butanol containing 4% paraformaldehyde (PFA) for 48 h. After washing four times in 80% *tert*-butanol for 10 min each, the samples were dehydrated in an ascending butanol series (90%, 96%, 3 × 100%) followed by three washes in pure acetone for 2 h each. Dehydrated samples were embedded in Technovit 8100 (Kulzer, Wehrheim, Germany), and semi-thin (8-µm) sections were prepared using an RM2245 rotary microtome (Leica Biosystems, Nußlock, Germany) with glass knives. The sections were alternatively mounted in two parallel series on microscope slides and hybridized with 500 nM of the general eubacterial probe EUB-338-Cy7 (5′-GCTGCCTCCCGTAGGAGT-3′) and a mix of four of the subsequent symbiont specific probes (Table S8) in 100 µL of hybridization buffer (0.9 M NaCl, 0.02 M Tris/HCl pH 8.0, 0.01% SDS) with 5 μg/ml DAPI for nuclear counterstaining. The sections were covered with a glass cover slip and incubated in a humid chamber at 50 °C overnight. The sections were washed in wash buffer (0.9 M NaCl, 0.02 M Tris/HCl pH 8.0, 0.01% SDS, 5 mM EDTA) at 50 °C for 1 h, followed by 30 min in water. After mounting the sections with Vectashield plus (Vector, Newark, CA, USA), they were imaged using a Leica THUNDER imager Cell Culture 3D equipped with a DFC9000GT monochrome camera (Leica, Wetzlar, Germany). The signals for the individual dyes/channels were recorded with the following LED8, filter cube (FC) and external filter wheel (EFW) settings: DAPI (5% 390nm LED, DFT51010 FC, 460/80nm EFW), background autofluorescence (50% 475nm LED, DFT51010 FC, no EFW), Rhodamine Green (50% 510nm LED, CYR71010 FC, 535/70nm EFW), Cyanine3 and Atto565 (50% 555nm LED, DFT51010 FC, 590/50nm EFW), Cyanine5 (50% 635nm LED, DFT51010 FC, 642/80nm EFW) and Cyanine7 (50% 747nm LED, CYR71010 FC, no EFW). Leica THUNDER images were processed in Leica Application Suite X software using the large-volume computational clearing algorithms with default settings.

## ACKNOWLEDGEMENTS

H.V., B.W., F.R., T.E. and M.K. express their gratitude to Henriette Ringys-Beckstein (Max Planck Institute for Chemical Ecology, Jena) for her technical support, and we thank Bruno Huettel (Max-Planck-Genome Center, Cologne) for his support in PacBio and Illumina genome sequencing. This research was supported by the Max Planck Society (H.V., B.W., F.R., T.E., M.K.), and the European Research Council through a Consolidator Grant (M.K., ERC CoG 819585 “SYMBeetle”). The authors thank Richard M. Twyman for professional editing of the manuscript.

## AUTHOR CONTRIBUTIONS

Heiko Vogel: Conceptualization, Supervision, Data curation, Formal analysis, Investigation, Methodology, Validation, Visualization, Writing – review and editing | Benjamin Weiss: Investigation, Validation, Visualization, Methodology | Fortesa Rama: Data curation, Formal analysis, Visualization | André Rinklef: Methodology | Tobias Engl: Investigation, Methodology | Martin Kaltenpoth: Formal analysis, Writing – review and editing, Investigation, Funding acquisition | Andreas Vilcinskas: Conceptualization, Project administration, Investigation, Supervision, Writing – original draft.

## COMPETING INTERESTS

The authors declare that they have no competing interests.

## DATA AVAILABILITY

The metagenomic data have been deposited in the European Nucleotide Archive (ENA) at EMBL-EBI under accession number PRJEB86762 (https://www.ebi.ac.uk/ena/browser/view/PRJEB86762). The bacterial genomic datasets for *Sulcia, Vidania, Purcelliella, Rickettsia, Wolbachia*, CAP and CPS, including genome annotation files, have been deposited in the Edmond data repository and can be accessed under the following weblink: https://doi.org/10.17617/3.4JXKRW. All other data are available from the corresponding author on reasonable request.

## ADDITIONAL FILES

Supplemental Material

Document S1 (Vogel-et-al_Supplement_S1.pdf). Figure S1. Supplemental figure on symbiont localization in *P. leporinus* tissue sections.

Document S2 (Vogel-et-al_Supplement_S2.xlsx). Tables S1-Sx. Supplemental methods (FISH probe sequences) and supplemental results on endosymbiont genome assemblies and annotations.

## Notes

https://www.ebi.ac.uk/ena/browser/view/PRJEB86762

https://doi.org/10.17617/3.4JXKRW

### Competing Interest Statement

The authors have declared no competing interest.

