## Supplemental File 1 for "A multi-partner symbiotic community inhabits the emerging pest *Pentastiridius leporinus*"

### SUPPLEMENTARY FIGURE S1

**Fig. S1. Details of bacterial localization in semi-thin transverse histological sections of *Pentastiridius leporinus* using fluorescent in situ hybridization (FISH).** The labeled eubacterial probe EUB-338-Cy7 (magenta) was used with DAPI to counterstain the host nuclei (cyan). **A.** Nerve tissue of the central ganglion with numerous *Arsenophonus* bacteria localized in the fat body (fb) and isolated cells identified in the somata (so). The neuropil (np) is largely free of bacteria. **B.** Numerous *Arsenophonus* bacteria in the fat body (fb) and in the muscles (m) of the *P. leporinus* thorax. **C-D.** *Arsenophonus* and *Wolbachia* cells were found to be localized in ovaries (ov) and fat body (fb) tissue. **E.** Bacteriome (symbiotic structure III), filled with high densities of unidentified bacteria (EUB-338) and numerous *Wolbachia* cells in parts of the fat body (fb). Tissue structure IV shows a high density of *Vidania* cells. **F-G.** Female reproduction-associated tissue with high densities of *Wolbachia, Arsenophonus* and partly intranuclear located *Rickettsia* cells. **H-I.** Detailed view of symbiotic structure IV with *Vidania* cells, symbiotic structure VI with densely packed *Sulcia* cells and fat body tissue with *Wolbachia* in a female *P. leporinus* specimen. The oocytes (oo) are free of bacteria. **J.** Symbiotic structure IV with *Vidania* in a female specimen. The spermatheca (sp) is free of bacteria. **K-L.** Isolated *Arsenophonus* cells identified between the testicular lobules of a male specimen. Otherwise the testicles (tes) are free of bacteria. *Wolbachia* cells can be found in parts of the fat body (fb). Scale bar = 50 µm.

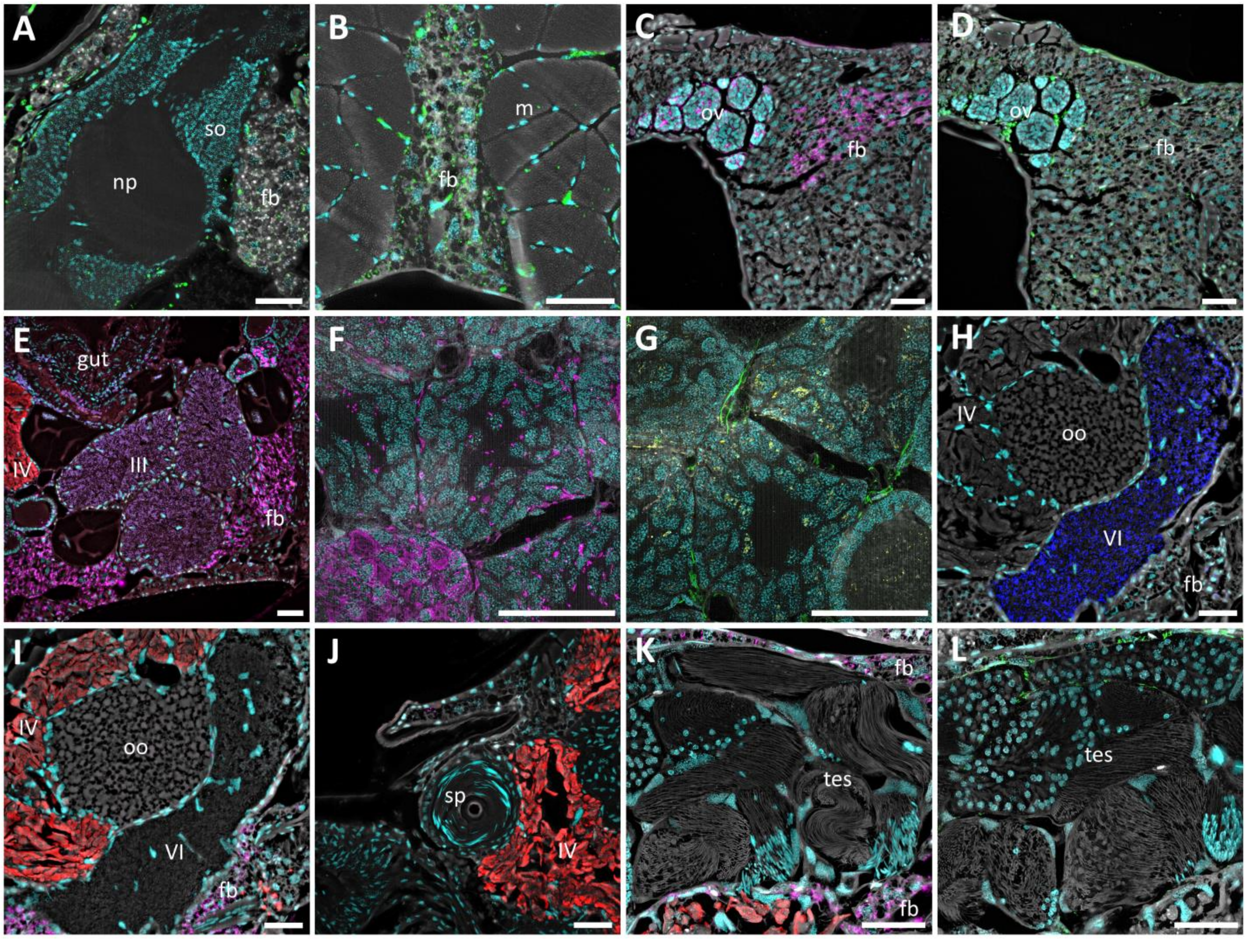
